# Truncation of IFT80 causes early embryonic loss in cattle

**DOI:** 10.1101/2021.07.02.450952

**Authors:** M. Sofía Ortega, Derek M. Bickhart, Kelsey N. Lockhart, Daniel J. Null, Jana L. Hutchison, Jennifer C. McClure, John B. Cole

## Abstract

Recessive alleles represent a risk in populations that have undergone bottleneck events. We present a comprehensive framework for identification and validation of these genetic defects, including haplotype-based detection, variant selection from sequence data, and validation using knockout embryos. Holstein haplotype 2 (HH2), which causes embryonic death, was used to demonstrate the approach. HH2 was identified using a deficiency-of-homozygotes approach and confirmed to negatively affect conception rate and stillbirths. Five carriers were present in a group of 183 sequenced Holstein bulls selected to maximize the coverage of unique haplotypes. Three variants concordant with haplotype calls were found in HH2: a high-priority frameshift mutation resulting in a deletion, and two low-priority variants (1 synonymous variant, 1 premature stop codon). The frameshift in intraflagellar protein 80 (*IFT80*) was confirmed in a separate group of Holsteins from the 1000 Bull Genomes Project that shared no animals with the discovery set. *IFT80*-null embryos were generated by truncating the *IFT80* transcript at exon 2 or 11 using a CRISPR-Cas9 system. Abattoir-derived oocytes were fertilized *in vitro* and embryos were injected at the one-cell stage either with CRISPR-Cas9 complex (n=100) or Cas9 mRNA (control, n=100) before return to culture, and replicated 3 times. *IFT80* is activated at the 8-cell stage, and IFT80-null embryos arrested at this stage of development, which is consistent with data from mouse hypomorphs and HH2 carrier-to-carrier matings. This frameshift in IFT80 on chromosome 1 at 107,172,615 bp (p.Leu381fs) disrupts WNT and hedgehog signaling, and is responsible for the death of homozygous embryos.

**Significance Statement:** Holstein haplotype 2 is an embryonic lethal present in 1.21% of the US Holstein cattle population, and unrecognized carrier-to-carrier matings are responsible for >$2 million/year in additional breeding expenses. A high-impact frameshift mutation in exon 11 of intraflagellar protein 80 (IFT80) was identified as the putative causal variant. Biallelic IFT80 knockout embryos were produced *in vitro* and compared to wild-type embryos. IFT80-null embryos consistently arrested at the 8-cell stage of development. The IFT80 protein expressed in knockout embryos had substantially altered protein structure, resulting in a loss of functional domains. These results validate the putative causal mutation observed in Holsteins. This system is a good model for investigating possible causal variants that affect livestock fertility early in development.

## Introduction

Known lethal recessive alleles account for substantial economic losses to cattle breeders (1), and 20 such defects (2) are routinely tracked in the US population of 9.4 million dairy cows. Economic losses resulting from embryonic death and stillbirths caused by these alleles have been estimated to cost farmers at least $11 million per year in the US (1). The increased use of artificial insemination on an already reduced genetic pool has exacerbated the problem by increasing the rate at which recessive lethal alleles can be spread in commercial herds. The HH2 allele, which accounts for annual fertility losses of more than $2 million in the US alone, is present in 1.66% of the domestic Holstein population (3) and has a significant negative effect on fertility in the form of early embryonic losses. The patrilineal nature of cattle reproduction and the availability of large numbers of genotyped animals provides an opportunity to identify and validate such recessive mutations and measure their potential impacts on other species.

## Results and Discussion

We first refined the HH2 genomic locus to narrow the window for variant discovery in subsequent sequencing analysis (Figure 1A). Our analysis was aided by the recent release of a high-quality reference assembly for taurine cattle (4) which had over 100-fold more contiguity than the previously used UMD3.1 reference (5). Using imputed SNP genotypes (6) derived from more than 3.7 million SNP genotypes of commercial cattle mapped to the new reference assembly, we were able to reduce the size of the locus from 8.9 Mbp to 1.5 Mbp. This reduced the number of candidate genes that were previously hypothesized to be causal for the lethality of the haplotype by 72%, leaving only seven genes in the refined window out of 25 original candidates.

**Figure 1.**
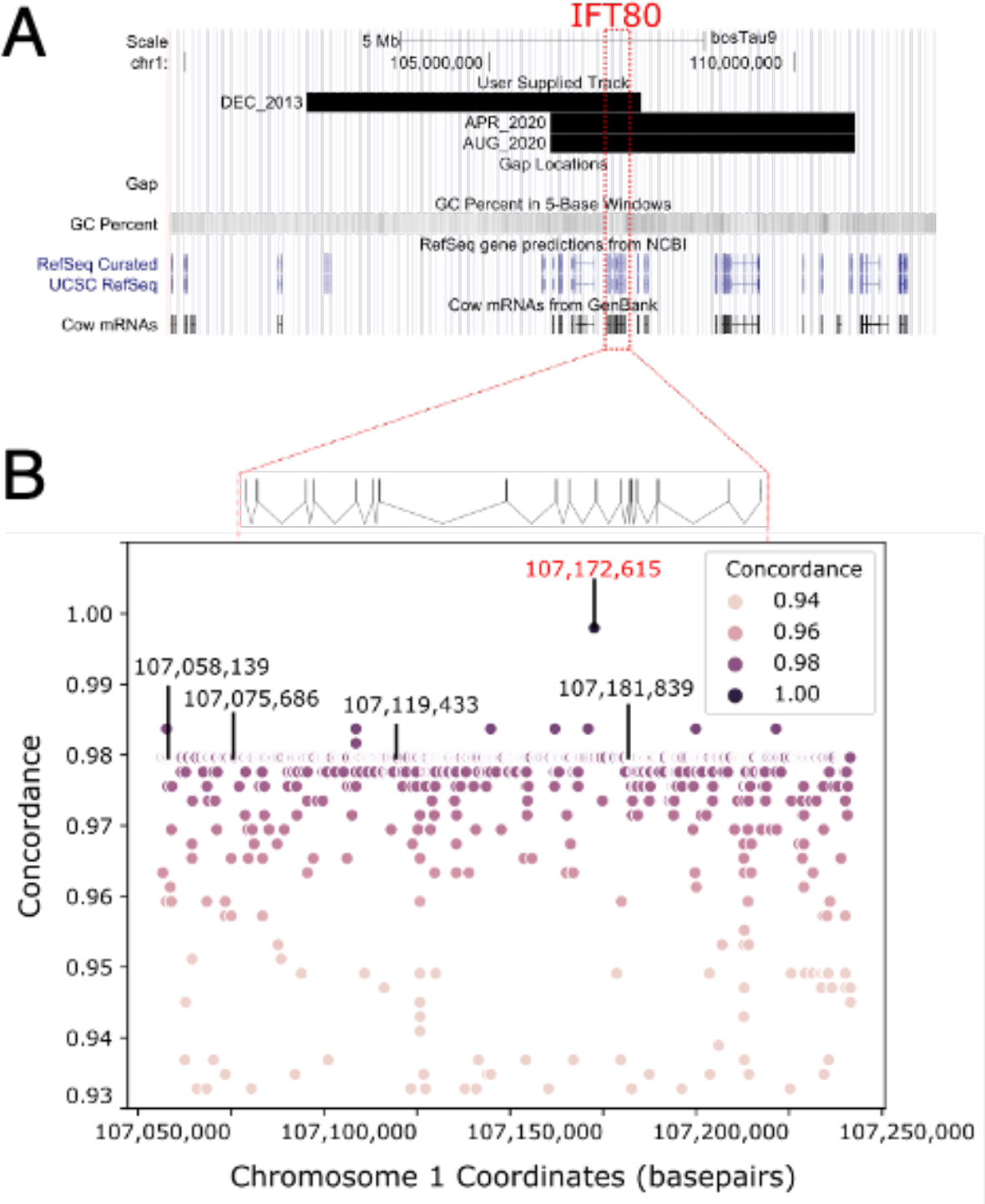
A UCSC genome browser plot (**A**) shows the locations of prior Holstein Haplotype 2 (HH2) loci as determined in December 2013, April 2020 and August 2020. The IFT80 gene is highlighted by red dashed lines. Concordance analysis of variant sites against expected carrier status (**B**) identified only one variant with > 99% concordance (highlighted in red). For genomic context, the IFT80 gene is drawn with exons (straight lines) and introns (diagonal lines) above the concordance plot. Only variants predicted to have high functional effect by SNPEff are highlighted by labels showing coordinate numbers.

Separate, non-overlapping datasets were used for causal variant discovery and validation. The discovery dataset included whole-genome sequence data for 9 carriers and 449 non-carriers from the Animal Genomics and Improvement Laboratory (Beltsville, MD), the Collaborative Dairy DNA Repository (Madison, WI) and the 1000 Bull Genomes Project (7). A concordance analysis was used to identify candidate causal variants within the haplotype locus, and one INDEL (Figure 1B) that was > 99% concordant with known carrier status was identified. One carrier animal was predicted to not have a heterozygous copy of the INDEL, but manual review of read pileups on the variant site identified the INDEL at a frequency of 25% which was below the threshold of initial detection (data not shown). We interpreted this result as complete concordance and attributed the lower proportion of variant reads to a bias in sampling the haplotype during sequencing library preparation. The concordant INDEL was located on btau1:107,172,615 and was found to induce a frameshift in the IFT80 gene’s 11th exon that led to early truncation of the protein product (Figure 2A). *In silico* translation of the expected protein product of the edited transcript suggested an early truncation of the IFT80 protein, resulting in a protein of 385 amino acids (49.5% of total) compared to the wild type (Figure 3D). The early termination of the gene interrupts one of seven WD 40-repeat containing domains (Pfam: SSF50978) and a predicted weak polyampholyte domain (https://mobidb.bio.unipd.it/A0A4W2EVR7), both of which likely impact the intra-flagellar transport of materials in the cell (8). Analysis of potential founders identified the bull Willowholme Mark Anthony (73HO0219, born 1975) as a likely candidate. Pedigree, SNP, and DNA sequence data show that he is the founder with respect to the SNP haplotype, but he is not a carrier of the actual IFT80 variant, which appears to have arisen in one of his daughters (Elysa Anthony Lea, HOCANF000003628269, born 1981) or grand-daughters (Comestar Laurie Sheik-ET, HOCANF000004425038, born 1986). Neither descendant is genotyped or sequenced and DNA is not available, so the true founder cannot be identified.

**Figure 2.**
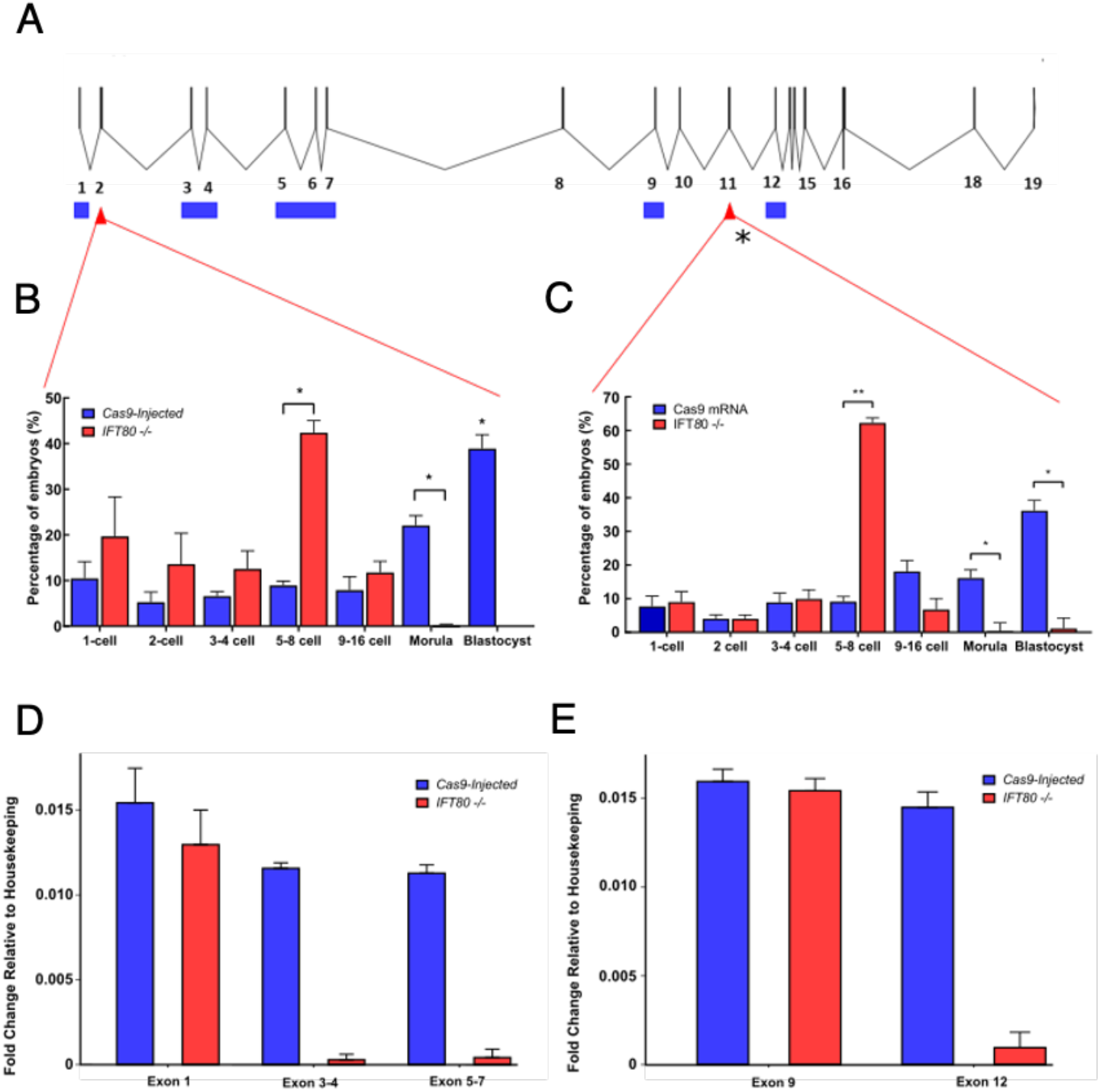
A diagram of the IFT80 gene (**A**) shows CRISPR-Cas9 edit sites (red triangles) and qPCR primer design sites (blue boxes). Gene exons are represented by vertical lines, whereas introns are represented by diagonal lines. Embryo counts for each developmental stage for exon 2 (**B**) and exon 3 (**C**) demonstrate that homozygous edit embryos (red) do not typically progress past the 8-cell stage compared to vector Cas9 controls (blue). Single (*) and double (**) asterisks indicate Student’s T test p-values that are less than 0.001 and 0.0001, respectively. Relative fold change of exons downstream of the exon 2 (**D**) and exon 11 (**E**) edit sites showed significant decreases in edited embryos (red) compared to the vector Cas9 controls (blue)

**Figure 3.**
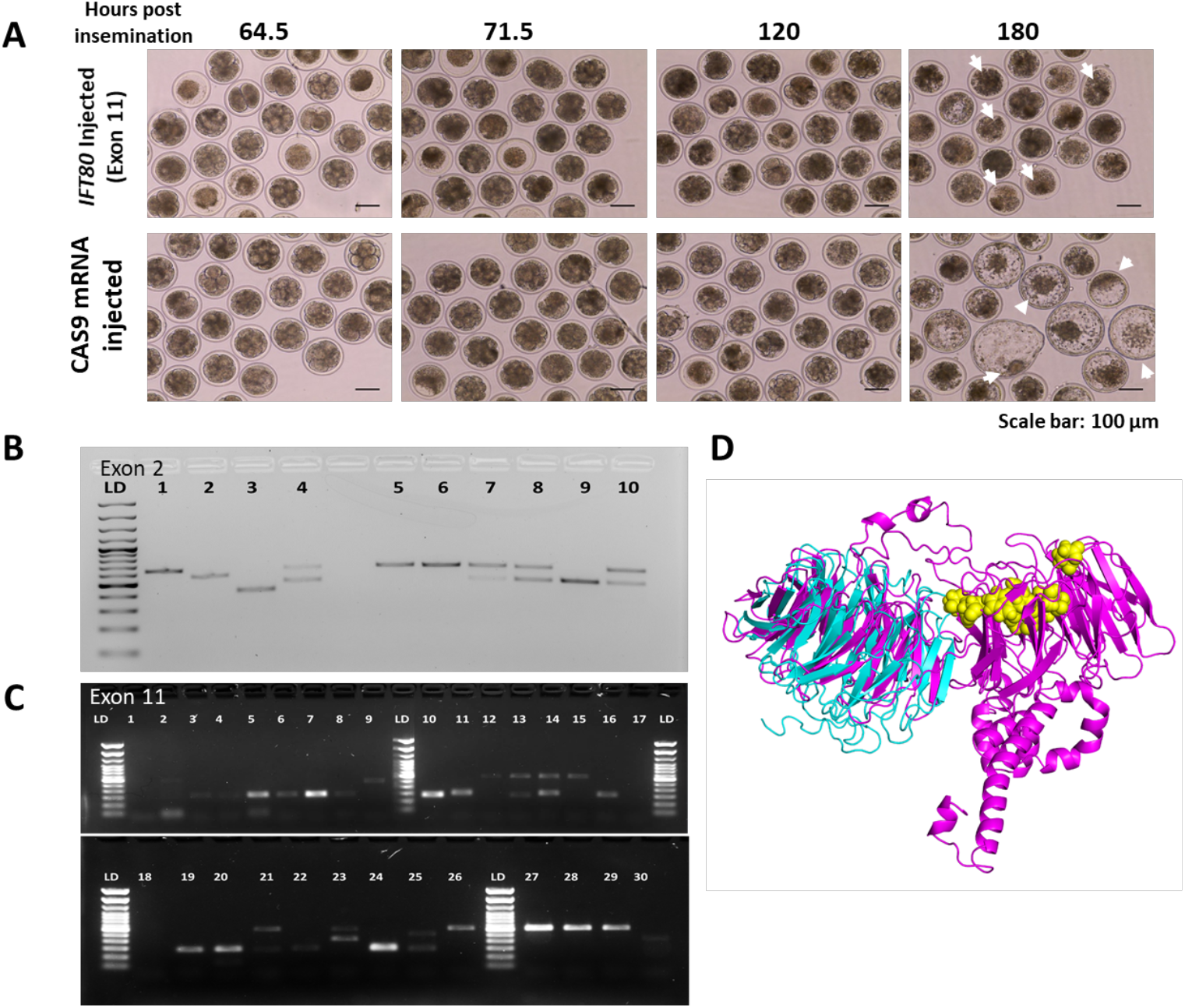
**A**. Development of control versus edited embryos. IFT80 injected group have increased number of degenerating embryos (white arrows,120 and 80 hours post insemination), after the 8-cell stage. In control embryos, development progressed normally with formation of blastocysts with compact inner cell mass and defined blastocoel (white arrows,180 hours post insemination). **B**. Embryo genotyping for exon 2 edits. Each numbered lane represents one embryo. Lanes 1, 5, and 6 show non-edited embryos (wild type); lanes 2, 3, and 9 show biallelic edits (absence of wild type band), and lanes 4, 7, 8, and 10 illustrate mono-allelic edits (both wild type, and edited band present). **C**. Embryo genotyping for exon 11 edits. Each numbered lane represents one embryo. Lanes 9, 12, 15, 26, 27, 28, and 29 show non-edited embryos (wild type); lanes 3 – 8, 10, 11, 16, 19, 20, 22, and 24 show biallelic edits (absence of wild type band), and lanes 2, 13, 14, 21, and 23 illustrate mono-allelic edits (both wild type, and edited band present). **D**. Protein model for IFT80 truncation at exon 11. The pink model represents the 777-amino acid wild-type protein, the blue model represents the 385-amino acid gene-edited protein (after truncation surrounding mutation), and the yellow globular regions indicate putative active binding sites.

We hypothesized that the early termination of the *IFT80* gene was the causal variant of the phenotype given its perfect concordance and the potential for the mutation to have a high impact on downstream protein function. To induce the phenotype, we used a CRISPR-Cas9 gene editing system (see Methods) to induce double-strand breaks at different exons at or upstream of the candidate causal variant site (Figure 2A). In the first model, IFT80 was truncated at exon 2, and in a second model, IFT80 was truncated at exon 11 (variant site). In both cases, treated embryos failed to progress past the 8- to 16-cell transition period (Figure 2A, 2c; P < 0.0001), and degenerated after this developmental stage (Figure 3A). As alternative splicing could potentially recover IFT80 protein function, we quantified the expression at different exons of IFT80 transcript. For embryos edited at exon 2, expression was measured at exons 1, 3–4 and 5–7 (before and after edit site; Figure 2D). For the embryos edited at exon 11 (exon with causal mutation), IFT80 expression was determined at exons 9–10 and 12 (before and after the edited exon; Figure 2E) by qPCR assays. Upstream regions of edited sites had no difference in IFT80 expression (Exon 1, P = 0.44; Exon 9-10, P = 0.62), between edited and non-edited embryos. Conversely, in both models (truncation at exon 2 or 11) downstream exons of edited site, had significantly lower IFT80 expression (P < 0.0001) in edited embryos than non-edited embryos, suggesting early termination of the transcript due to the mutation or the altered splicing of these exons from the transcript (Figure 2D). Differential expression was most pronounced in exon 12, in which the edited cell line had 62.5% lower fold expression of the exon than the wildtype (Figure 2E).

Mutations to the *IFT80* gene are linked to asphyxiating thoracic dystrophy 2 (ATD2) disease in human (9) and hypomorphic expression of the gene has been shown to result in low postnatal survival in gene-trap mice (10). Previous surveys have speculated as to when IFT80 gene expression is essential during development (10), and our results suggest that expression is needed to progress past the 8- to 16-cell developmental phase. This is consistent with previous RNA surveys of cattle preimplantation embryonic development (11), and with the known fertility effects of HH2 in cattle (1, 3). The results of editing experiments suggest that the carboxy-terminal WD 40-repeat or the polyampholyte domains are essential for embryonic development. Recent studies suggest that deletion of IFT80 disrupt FGF2 signaling pathway, affecting cell proliferation, and self-renewal of odontoblasts (12). In the preimplantation mammalian embryo, FGF2 signaling is essential for lineage commitment and blastocyst formation (13, 14), in which disruption of FGF2 signaling coincides with the developmental arrest observed in this study.

Using genotypes from millions of Holstein cattle and CRISPR-Cas9 gene editing, we were able to track and validate a recessive lethal allele in the IFT80 gene, respectively. This approach conclusively identified the target mutation in such a way that it provided further insight into the function of the gene in development of other mammalian species. There is a further economic benefit to this discovery, as identification of the exact causal mutation underlying this haplotype will benefit dairy producers by allowing them to avoid carrier-to-carrier matings which result in pregnancy losses responsible for ∼$437,000/year in US Holsteins. Concordance analysis and molecular validation on naturally occurring genetic variants represent a potent tool for ongoing efforts to translate genotype to phenotype in the field of quantitative genetics.

## Materials and Methods

### Haplotype discovery

Possible autosomal recessive mutations are routinely tracked during the imputation step of the national genomic evaluations calculated by the Council on Dairy Cattle Breeding (CDCB; Bowie, MD). Genotypes from 48 different arrays are currently stored in the CDCB database, with SNP counts ranging from 2,900 to 777,962, and are imputed to a set of 78,964 SNP (80k) for genomic evaluation. No single chip includes all these markers, regardless of density, and genomic calculations require that genotypes are available for all 80k SNP. Chromosomes are phased into maternal and paternal haplotypes and the genome is divided into 100-SNP segments. Inheritance of those segments is traced through the population and missing SNP are filled in using a segment from the pedigree when available, and using the most common segment in the population when pedigree information is not available (3). Most males have complete pedigrees available for many generations, but many females do not. For this study, haplotyping was performed using the program Findhap version 3 (15). Haplotypes that either never appear in the homozygous state, or which appear with far lower frequency than expected, are flagged as possible recessives (3). Such haplotypes are then tested for effects on fertility and stillbirth. Founder animals are identified from the pedigree when possible. Currently, 26 such conditions are tracked in the U.S. dairy cattle population (20 genetic diseases and 6 coat-color and polled variants), and causal mutations are known for all but the HH2 haplotype.

In April of 2011 a haplotype affecting fertility was identified on *Bos taurus* autosome (BTA) 1 in the region of 92–97 Mbp and designated as HH2 (Figure 1A). At that time, a second haplotype on BTA 1 at 101–106 Mbp with a similar carrier list, lack of homozygotes and negative effects on fertility also was identified. When a new SNP list was introduced in August 2011 the haplotype in the 101–106 Mbp region was no longer flagged as a potential lethal. A new 61,000 SNP list was introduced in December 2013, and the haplotype segment upstream from HH2 again appeared on the list of possible lethal haplotypes. In April 2020 the location of HH2 was changed to correspond with the upstream segment based on results from fine-mapping described below. Haplotype blocks are formed using fixed numbers of SNP, rather than LD boundaries, so it can sometimes be difficult to narrow the candidates down to a single haplotype. In this case, both segments appear to track the causal variant with high concordance.

### Whole-genome sequencing, variant calling, and annotation

The dataset used for causal variant discovery and validation included whole-genome sequence data for 9 carriers and 763 non-carriers, including animals from the Ayrshire, Holstein, and Jersey breeds sequenced by the Animal Genomics and Improvement Laboratory (Beltsville, MD), the Collaborative Dairy DNA Repository (Madison, WI), and the 1000 Bull Genomes Project (7). Sequencing depth for these animals averaged 14.7x. Sequence data were aligned to the ARS-UCD1.2 reference genome (4) using BWA-MEM as implemented in bwa version 0.7.15-r1140 (16) and converted to the binary Sequence Alignment/Map format by samtools version 1.3 (17). 1000 Bull Genomes run8 data were processed using version 4 of the Genome Analysis Toolkit (18), and included genotyping, variant recalibration for SNP and INDEL, removal of individuals that failed QC, and filtering of monomorphic alleles. The resulting VCF file was filtered to remove reads with QS < 10 and those identified as “LOWQUAL”. Sequence data variants were identified using the Samtools v1.9 and BCFtools v1.9 (19) and reads passing edits were aligned against version ARS-UCD1.2 of the bovine genome (4). The list of filtered variants was annotated with the SNPEff (20) utility using the Ensembl annotation of the cattle reference genome (version 101). Variants listed as “High”, “Medium” and “Low” priority within the haplotype region by SNPEff were selected for further analysis. Impacts on predicted protein translation were identified using custom scripts.

### Production of IFT80^-/-^ embryos

#### Guide RNA design

Guide RNAs (gRNAs) were designed to target exon 2 or exon 11 (Table 1), using the GPP portal from the Broad Institute (https://portals.broadinstitute.org/gpp/public/analysis-tools/sgrna-design). To select gRNAs, two criteria were used: (1) those with the highest on-target score after design based on the GPP portal and the guide RNA checker portal (IDT technologies, San José, CA), and (2) those having the least similarity with other bovine sequences containing a protospacer adjacent motif (PAM). Selected gRNAs, as well as the universal 67mer tracrRNA® for form guide complexes were ordered from IDT technologies.

**Table 1.**
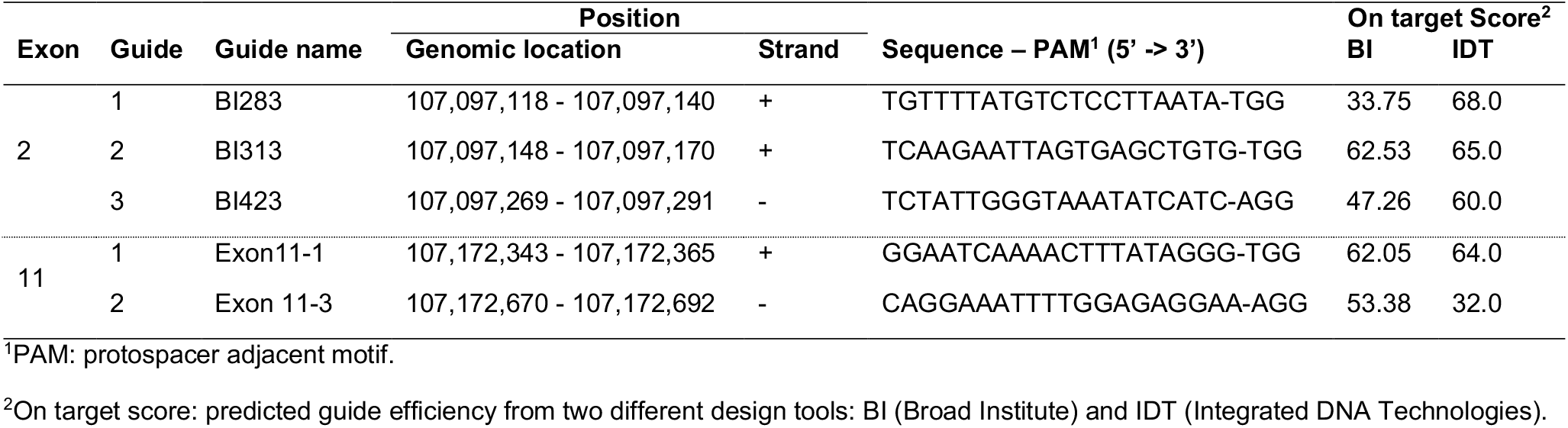
IFT80 guide RNA design table for exons 2 and 11.

The gRNAs and tracrRNA were resuspended to a concentration of 500 ng/μl. Each guide was then annealed with equal amounts of tracrRNA by heating at 95 °C for 5 min and then cooled down from 95 to 25 °C using a gradient of −1 °C/12 s in a Veriti 96-well gradient thermal cycler (ThermoFisher, CA) to obtain a concentration of 250 ng/μl gRNA-tracrRNA complex. Prior to zygote injection, gRNA-tracrRNA complexes were annealed to CAS9 mRNA (IDT Technologies) by incubation at room temperature for 15 min. The final injection solution was 50 ng/μl each gRNAs complex and 20 ng/μl Cas9 mRNA. For each exon targeted, guides were injected in pairs, to create a sizeable cut in the sequence and facilitate rapid genotyping on an agarose gel. The expected cuts for guides targeting exon 2 was 82 bp and for those targeting exon 11 were expected to generate a cut of 331 bp.

#### Production of embryos in vitro and zygote microinjection

Embryos were produced in vitro with a single sire known to be of high fertility in vitro as previously described (21). At the end of fertilization, putative zygotes (oocytes exposed to sperm) were denuded from the surrounding cumulus cells and split in two groups: injected only with Cas9 mRNA and injected with gRNA/Cas9 mRNA against IFT80. During the injection procedure zygotes were maintained manipulation medium (22), and then transferred to four-well dishes in groups of up to 50 zygotes in 500 µl of SOF-BE2 (23), covered with 300 µl of mineral oil per well at 38.5 °C in a humidified atmosphere of 5% (v/v) O2 and 5% (v/v) CO2. Percentage of putative zygotes that cleaved was determined at day 3 of development (day 0 = day of insemination) and blastocyst rate was estimated at day 8 of development. In addition, to determine any developmental arrest, cell stage of all embryos was recorded at days 64.5, 71.5, 120 and 180 h of development (Figure 2B, 2C and 3A).

#### Editing efficiency

To determine editing efficiency of the guides, embryos subjected to microinjection were collected individually at day 8 of culture, placed in 6 µL of Embryo Lysis Buffer (22), and subjected to 30 min at 60°C, followed by 10 min at 85 °C in a Veriti 96-well gradient thermal cycler (ThermoFisher, CA). Lysed embryo solution was used as template DNA.

For genotyping, the region surrounding the target sequenced of the gRNAs was amplified by endpoint PCR using the following primers (5’→3’): exon 2 F-GGTTTCTTATCCTGCTTTCCATTC; R-GAAATTGAGTGTGAACCTTGGG, and for Exon 11 F-CACTGTTTAGGACTCTGCCT; R-CTCTCTGAGTAATGATACCATAGCA. The PCR products were 639-and 620 bp for exon 2 (Figure 3B) and 11 (Figure 3C), respectively. The PCR reaction, amplification (annealing temperature 53.4 °C), and visualization of bands was performed as previously described (22). Editing efficiency was calculated from 3 separate in vitro embryo production runs, with 100 embryos per treatment group per replicate. Editing efficiency for exon 2 was 26% bi-allelic, 32 % mono-allelic, and 42% non-edited, and for exon 11 was 50% bi-allelic, 23% mono-allelic, and 27% non-edited.

#### Nucleic acids isolation and RT-qPCR expression analysis

DNA and RNA were isolated from individual 8-cell embryos (65–72 h post insemination) injected against IFT80 or injected only with Cas9 mRNA. Embryos were collected and lysed individually in 25 µl of RLT Lysis buffer (Qiagen, Valencia, CA), lightly vortexed, transferred to a nano column (PuroSPIN™ Luna NANOTECH, Toronto, ON, CA), and centrifuged for 30 s (all centrifugations were performed at 16,000 g). At this point, DNA is trapped in the column and the flow-through contained the RNA. The column was placed in a new collection tube and stored at room temp for later DNA purification. The flow-through containing RNA was mixed with one volume of 70% ethanol, transferred to a new nano column, and centrifuged for 30 s. Immediately after, 600 µl of buffer RW1 (Qiagen) were added to the column, and centrifuged for 30 s. Flow-through was discarded from the collection tube and 500 µl of buffer RPE (Qiagen) were added to the column and centrifuged for 30 s. After discarding the flow-through, 500 µl of 80% ethanol were added to the column, and centrifuged for 2 min. To elute RNA, the column was placed in a new microcentrifuge tube, and 15 µl of RNAse-free water were added directly to the membrane of the column, incubated at room temperature for one minute and finally centrifuged for 1 min. To complete DNA isolation, 500 µl of 70% ethanol were added to the nano column containing DNA, followed by a 1-min centrifugation. Then, the column was placed in a new microcentrifuge tube and 15 µl of EB buffer (Qiagen) preheated at 70 °C were added directly to the membrane of the column, incubated at room temperature for 10 min, followed by 1 min centrifugation to elute DNA. Embryos edit status was first assessed using PCR across the site of the edit as described above, and the RNA from homozygous edited embryo cells were selected for further analysis. Due to the small amount of total RNA in each cell, homozygous embryos were pooled in groups of four prior to cDNA creation. Three pools of edited and control embryos were analyzed in subsequent analyses. Two strand cDNA synthesis was conducted using the high-capacity cDNA Reverse transcription kit (ThermoFisher) reverse transcriptase following manufacturer instructions. Primers sequences for each locus were designed using the PrimerQuest™ (IDT technologies). All primers (Table 2) were validated following procedures previously described (24). qPCR was conducted using the CFX384 Touch Real-Time System (Bio-Rad) using 10 µl of reaction mix as previously described (22). The gene SDHA was chosen as a suitable housekeeping gene for relative expression calculation in bovine embryos (25). Fold changes were calculated relative to the housekeeping gene (2ΔCT) as previously described (22, 24).

**Table 2.**
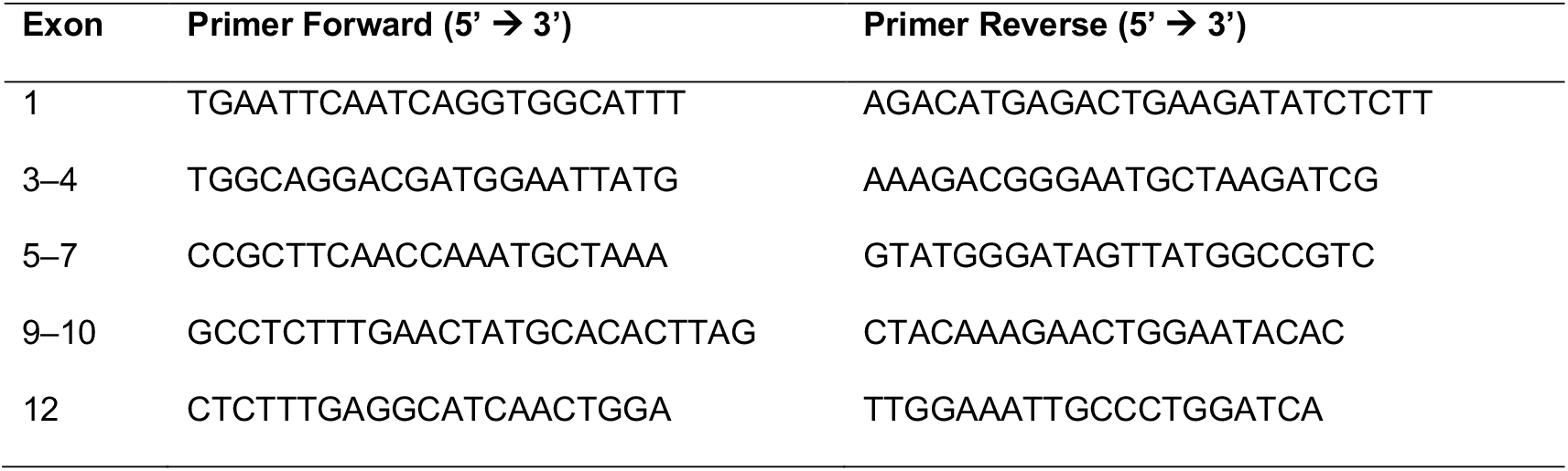
Primer sequences for gene expression analysis.

### Statistics

Differences in gene expression and development were analyzed by least-squares analysis of variance of the ΔCT values using the GLM procedure of the Statistical Analysis System version 9.4 (SAS Institute Inc., Cary, NC, USA). Replicate and treatment were included as main effects in the model.

## Data Availability

### Performance and Pedigree Data

The performance and pedigree data used to compute HH2 effects are controlled by the Council on Dairy Cattle Breeding (CDCB; Bowie, MD). Requests to access CDCB data may be sent to João Dürr, CDCB Chief Executive Officer.

### Genotype and Whole-Genome DNA Sequence Data

Access to SNP genotype data and whole-genome DNA sequence data for bulls held by the Collaborative Dairy DNA Repository (CDDR; Verona, WI) must be requested from Jay Weiker, CDDR Administrator.

### 1000 Bull Genomes Data

Access to whole-genome DNA sequence data from bulls distributed in Run7 is restricted to project members. Inquiries should be directed to Dr. Ben J. Hayes.

## Acknowledgments

Bickhart was supported by appropriated projects 8042-31000-001-00-D, “Enhancing Genetic Merit of Ruminants Through Improved Genome Assembly, Annotation, and Selection” and 5090-31000-026-00-D, “Investigating Microbial, Digestive, and Animal Factors to Increase Dairy Cow Performance and Nutrient Use Efficiency”, of the Agricultural Research Service (ARS) of the United States Department of Agriculture (USDA). Cole, Hutchison, and Null were supported by appropriated project 8042-31000-002-00-D, “Improving Dairy Animals by Increasing Accuracy of Genomic Prediction, Evaluating New Traits, and Redefining Selection Goals”, of ARS, USDA. McClure was supported by appropriated project 5090-31000-026-00-D, “Investigating Microbial, Digestive, and Animal Factors to Increase Dairy Cow Performance and Nutrient Use Efficiency” of ARS, USDA. Clark was supported by USDA NIFA Grant 2019-38420-28972.

The authors thank US dairy producers for providing phenotypic, genomic, and pedigree data through the Council on Dairy Cattle Breeding under USDA Agricultural Research Service (ARS) Material Transfer Research Agreement 58-8042-8-007. Access to whole-genome sequence data from the Collaborative Dairy DNA Repository (Madison, WI) was provided under USDA-ARS Material Transfer Research Agreement 58-8042-9-0010F. Access to whole-genome sequence data from the 1000 Bull Genomes Project was provided under USDA-ARS Material Transfer Agreement 14358. The authors also thank Bethany Bauer and Joshua Benne from the University of Missouri for their assistance with microinjections of bovine embryos.

Mention of trade names or commercial products in this article is solely for the purpose of providing specific information and does not imply recommendation or endorsement by the US Department of Agriculture. The USDA is an equal opportunity provider and employer.

## Notes

**Competing Interest Statement:** The authors declare no competing interests.

### Competing Interest Statement

The authors have declared no competing interest.

### Summary of Updates

Rewrote abstract. Expanded and revised Materials & Methods. Updated figures.

